## Supplementary Figures for "Broadly neutralizing SARS-CoV-2 antibodies through epitope-based selection from convalescent patients"

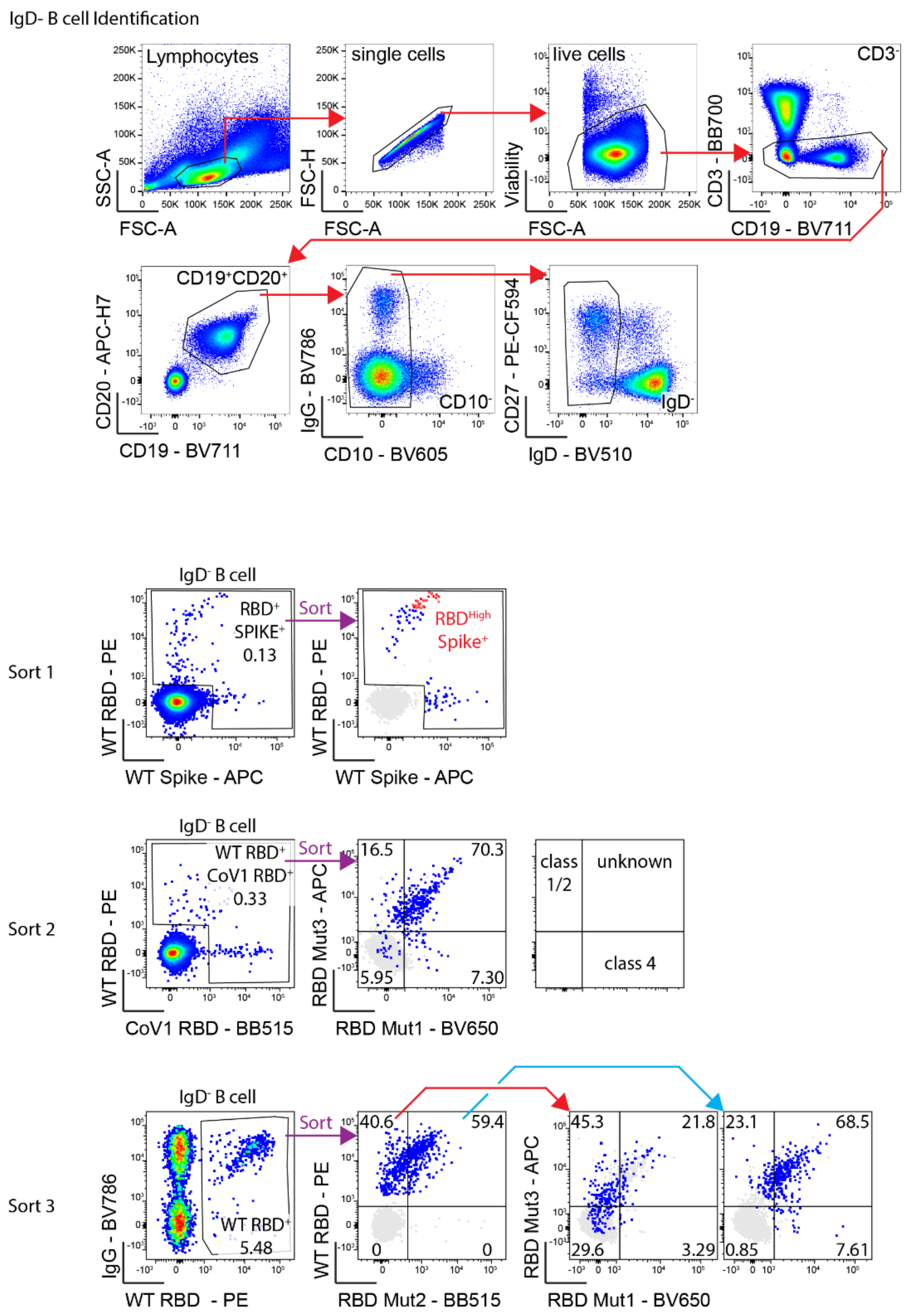


**Supplementary Fig. 1: Overview of the PBMC sorting strategy.** Representative flow cytometric plots of circulating IgD- CD27+ memory B cells in a convalescent patient one month after SARS-CoV-2 infection. The cells were also stained with fluorescent tetramers of SARS-CoV-2 RBD and SARS-CoV-2 spike protein for the Sort 1 strategy. For the Sort 2 and Sort 3 strategies, cells were strained with WT RBD, Mut1, Mut2 and SARS-CoV-1 RBD (replaced by Mut3 RBD in Sort 3).

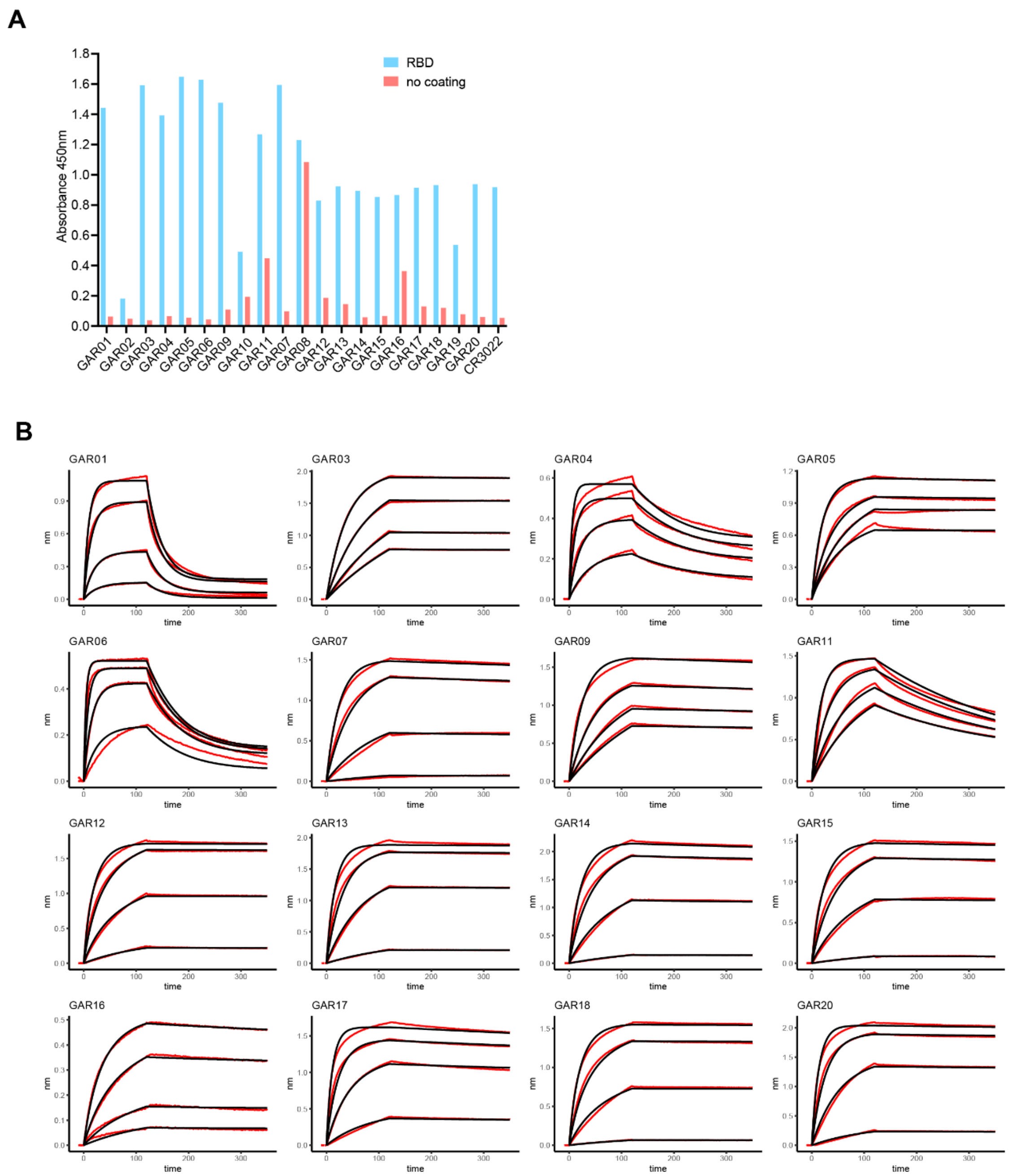


**Supplementary Fig. 2: Binding of monoclonal antibodies to SARS-CoV-2 RBD.** A: Purified antibodies were initially screened by ELISA for their binding to the RBD. RBD was coated on a Maxisorp ELISA plate at 2 µg/ml, incubated first with the purified antibody at 100 µg/ml and then with HRP conjugated to anti-human IgG. B: Affinity measurements by BLI. Streptavidin sensors were coated with biotinylated antibodies at 40 µg/ml, and association and dissociation constants measured using 2-fold serial dilutions of the SARS-CoV-2 RBD from

400 to 50 nM. Raw binding curves are represented in red and the fitted curves in black ($K_D$ values in Figure 1C).

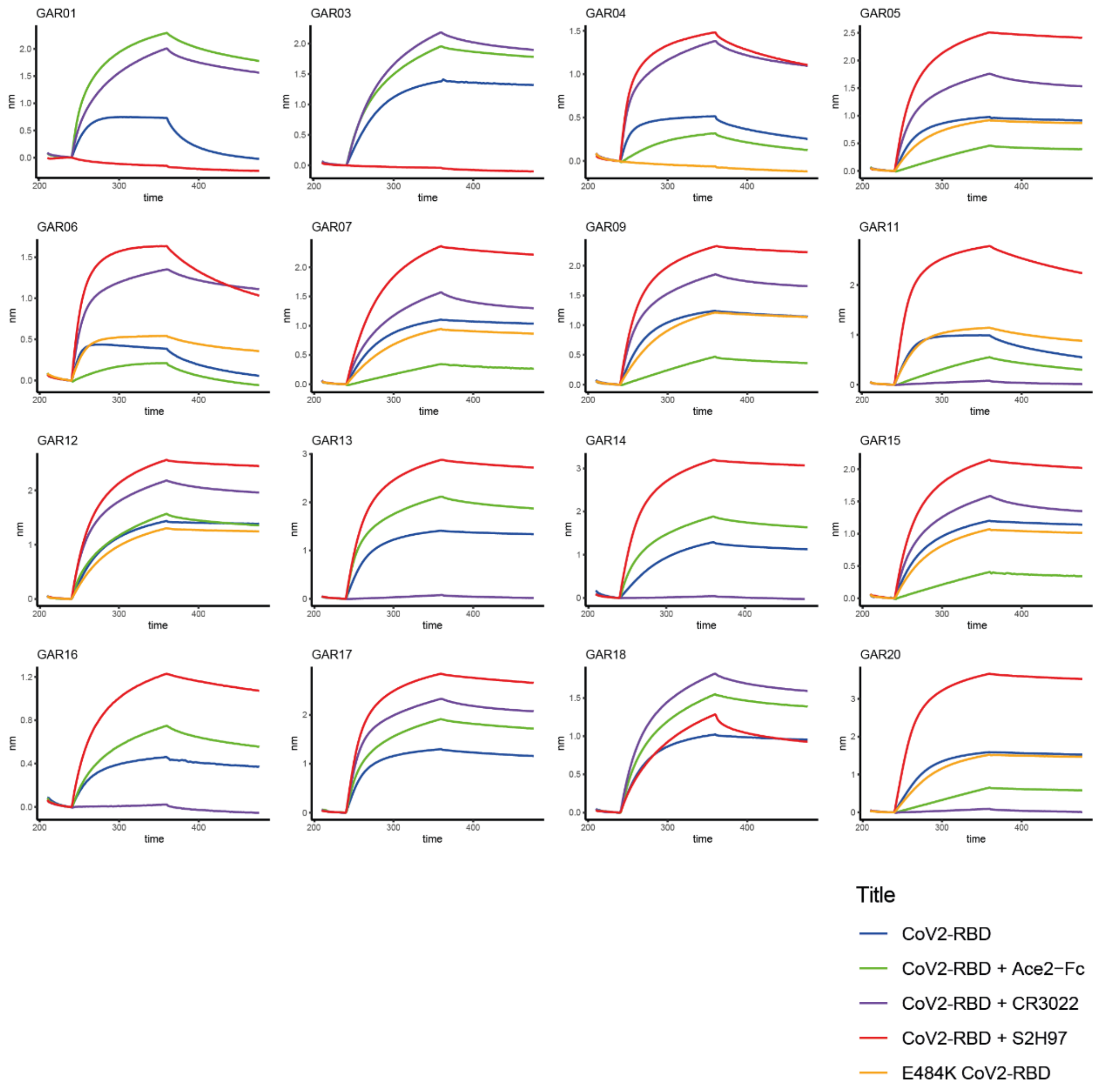


**Supplementary Fig. 3: Epitope mapping through BLI competition.** Antibodies were initially assessed for binding to the RBD (500 nM) pre-incubated with human ACE2 Fc (1 µM) to identify class 1 and 2 antibodies. Additive binding signal indicates co-binding while a decrease or absence of signal indicates competition. Non-class 1 and 2 antibodies were then evaluated for binding to RBD (500 nM) pre-incubated with CR3022 IgG or with S2H97 IgG (1 µM) to identify the class 4 or 5 antibodies, respectively. Finally, class 1/2 antibodies were evaluated for binding to the RBD E484K mutant (500 nM) to suggest class 2 antibodies.

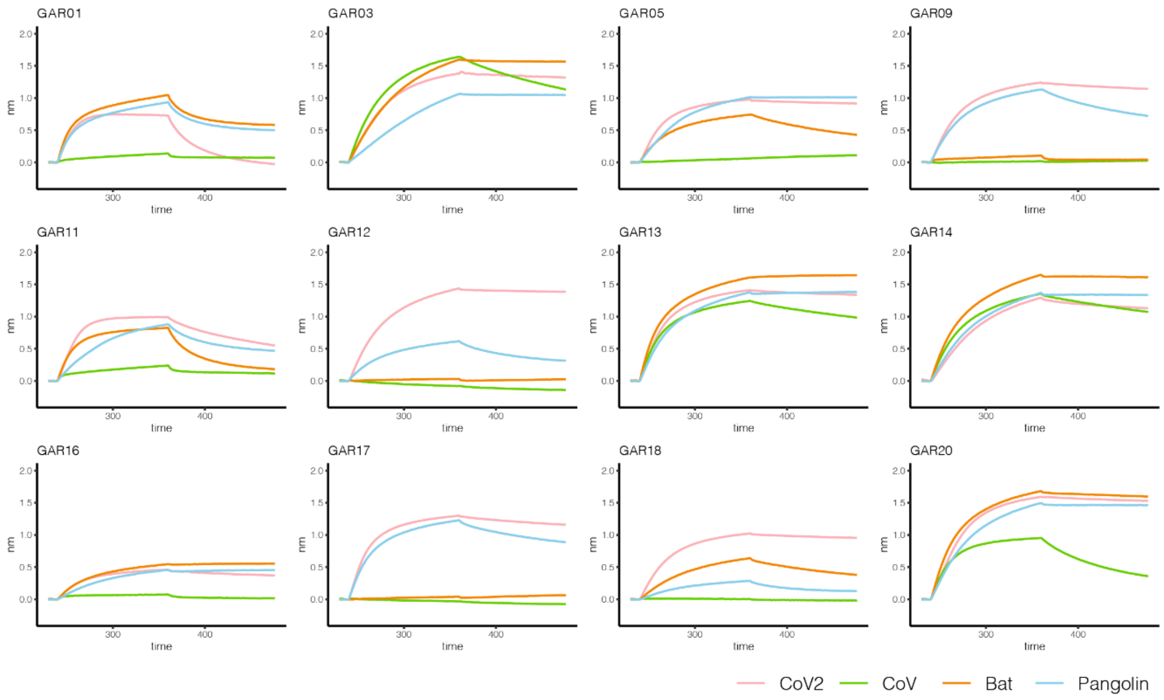


**Supplementary Fig. 4: Cross reactivity to other sarbecovirus RBDs.** Monoclonal antibodies were evaluated for binding to SARS-CoV-1, Bat RaTG13 and Pangolin CoV by BLI. Biotinylated antibodies were captured onto streptavidin sensors and incubated with the RBDs at 500 nM.

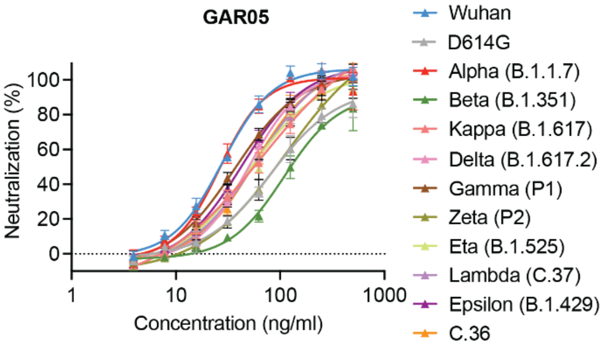


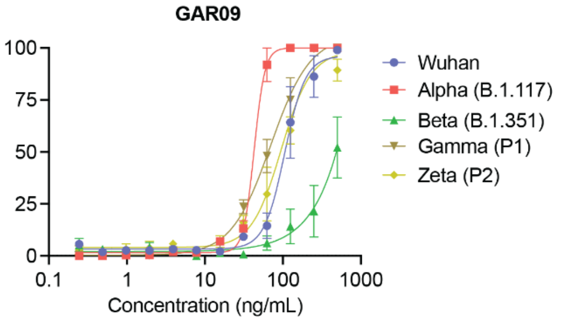


| strain | Wuhan | D614G | Alpha | Beta | Kappa | Delta | Gamma | Zeta | Eta | Lambda | Epsilon | C.36 |
|---|---|---|---|---|---|---|---|---|---|---|---|---|
| $IC_{50}$ (ng/ml) | 27.8 | 82.0 | 26.3 | 109.1 | 65.8 | 50.1 | 36.6 | 114.7 | 50.5 | 59.6 | 43.0 | 56.0 |

| Wuhan | Alpha | Beta | Gamma | Zeta |
|---|---|---|---|---|
| 106.0 | 42.8 | >10000 | 69.7 | 95.5 |

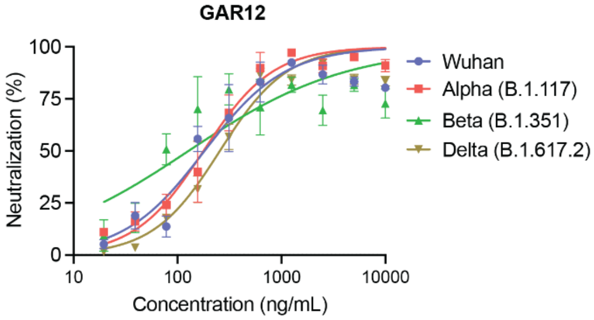


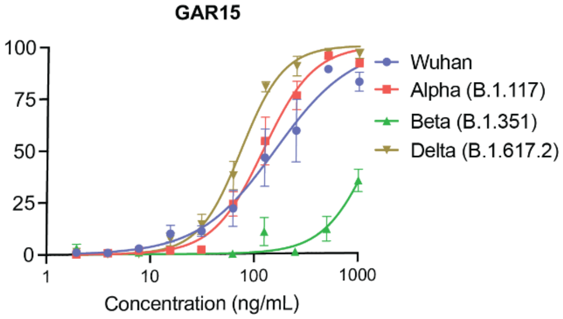


| Wuhan | Alpha | Beta | Delta |
|---|---|---|---|
| 181.9 | 178.0 | 127.6 | 255.5 |

| Wuhan | Alpha | Beta | Delta |
|---|---|---|---|
| 157.2 | 120.9 | 1403 | 73.8 |

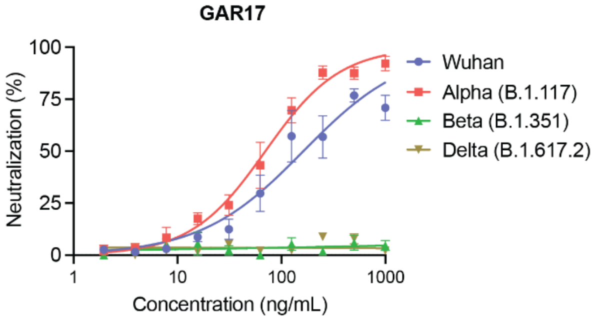


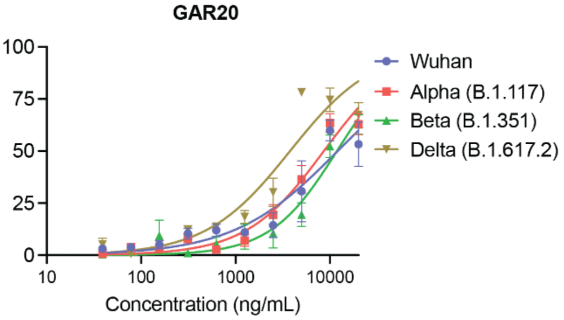


| Wuhan | Alpha | Beta | Delta |
|---|---|---|---|
| 162.5 | 70.0 | nn | nn |

| Wuhan | Alpha | Beta | Delta |
|---|---|---|---|
| 12044 | 8601 | 11828 | 3772 |

**Supplementary Fig. 5: Neutralization of SARS-CoV-2 VOCs.** Antibodies were evaluated for *in vitro* live SARS-CoV-2 virus neutralization in Vero E6 cells or HEK293T cells expressing hACE2 (for the Delta variant only) (n=4).

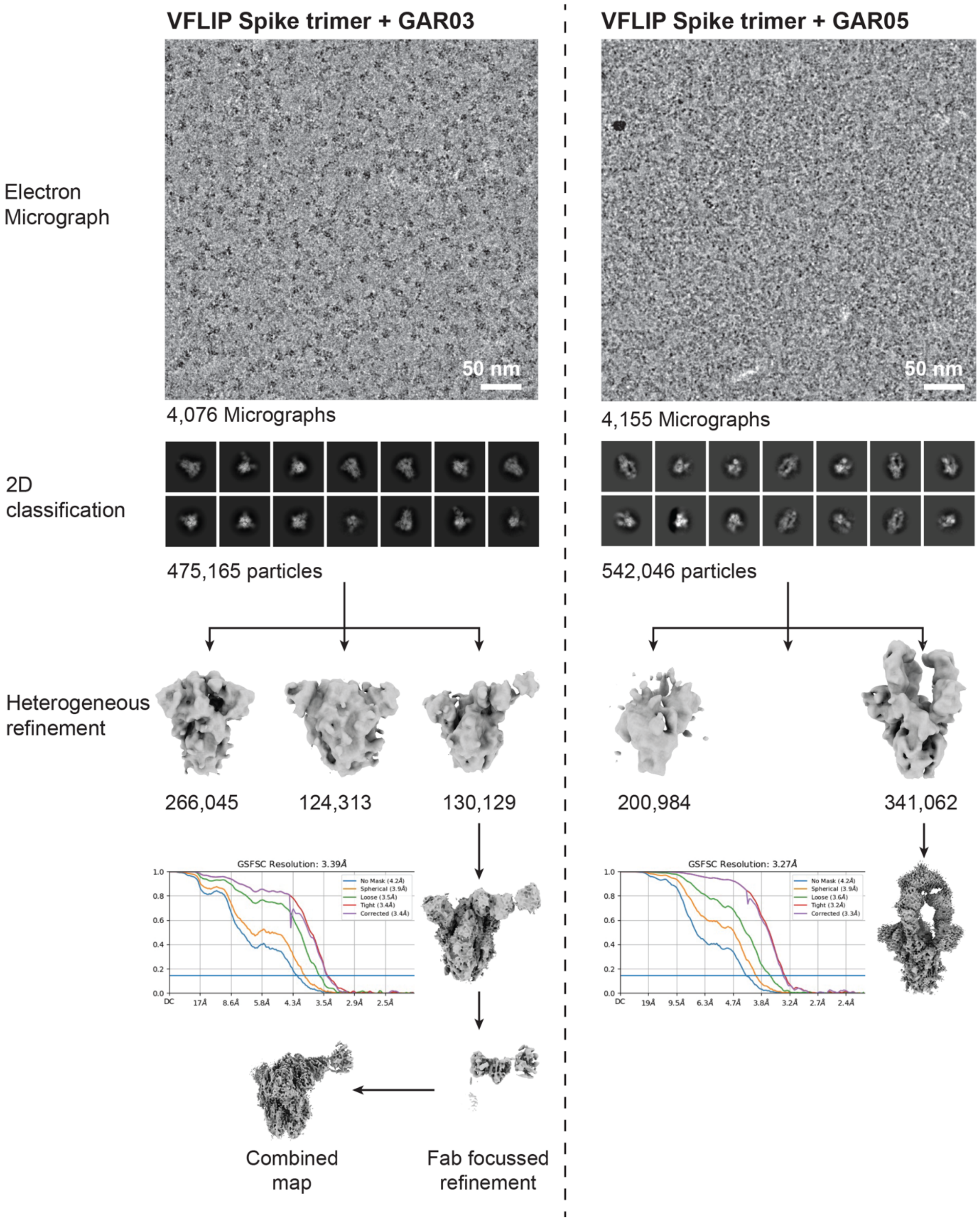


**Supplementary Fig. 6: Cryo-EM data processing flowchart.** Cryo-EM data processing strategy along with the Fourier shell correction (FSC) curves on the final maps.

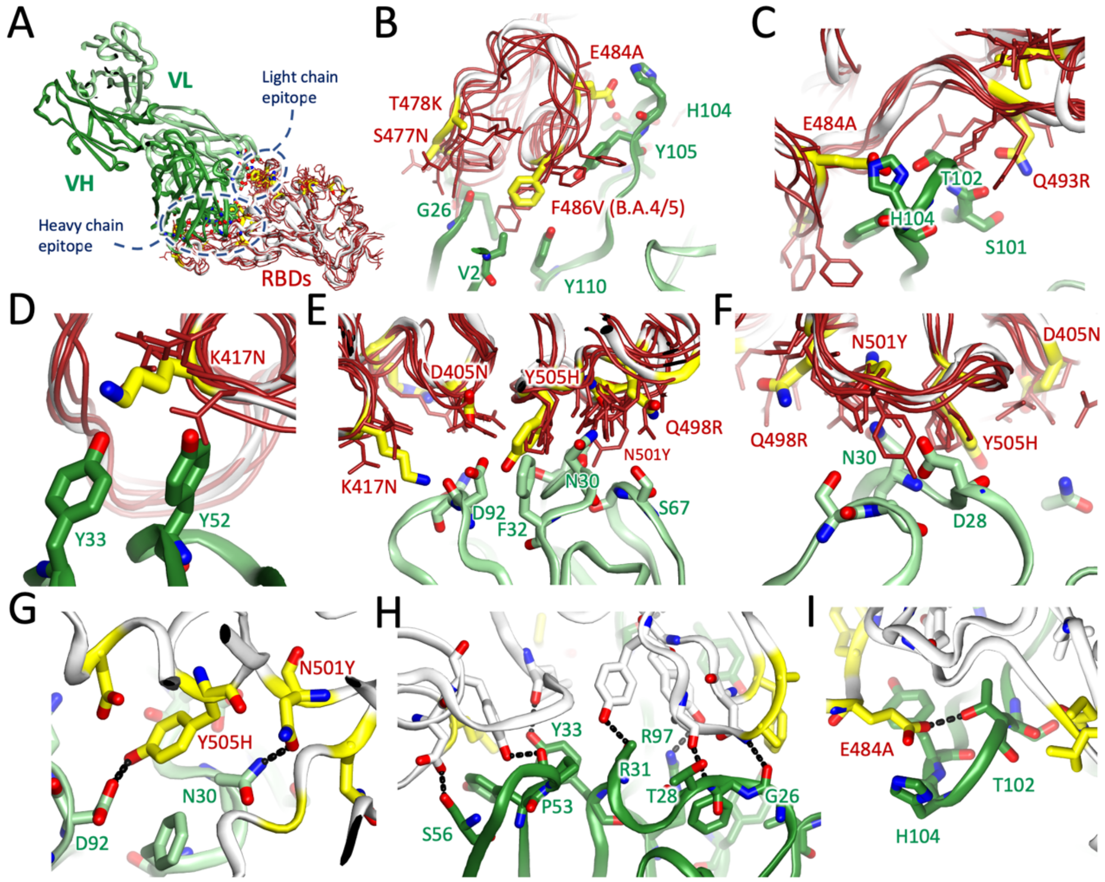


**Supplementary Fig. 7: GAR05-RBD superposed with omicron lineage structures**. A: GAR05 heavy and light chain (dark and pale green cartoons) bound to Wuhan RBD (cream cartoon) superposed with omicron lineage RBDs (thin red cartoon and sticks). Omicron structures include those from lineages B1.1.529, BA.2, and BA.4/5, and come from a variety of conformational contexts*. Residues shown as yellow sticks mark positions mutated in omicron lineages. The epitopes recognised by GAR05 heavy and light chains are indicated. The perspective is equivalent to that shown in Fig. 2B) Close-up views of GAR05 heavy chain residues in proximity to omicron variant positions. Panel B highlights several mutations (S477N, T478K, E484A and F486V (exclusive to B.A.4/5)), which adorn a relatively flexible loop comprising one end of the saddle to which ACE2 binds.  Panel C focuses on the E484A and Q493R positions, whilst panel D highlights the K417N position. Panels E and F highlight light chain residues in proximity to omicron mutations. Panels G-I detail hydrogen bonds (black dashed lines) between GAR05 and the RBD for the light chain (panel G) and the heavy chain (panels H and I). The bulk of these involve RBD side chains or main-chain residues not mutated in omicron variants (RBD features coloured cream), with notable exceptions of the light-chain centric Y505H and N501Y (coloured yellow, panel G) and heavy-chain centric E484A (panel I)

*Omicron structures shown as thin red wire cartoons and sticks comprise; B.1.1.529 in the up and down conformations (PDB 7tgw chains A and B, at 3.0 Å [1]), B.1.1.529 in the exclusively down conformation (PDB 7wp9, chain A, at 2.56 Å [2]), B.1.1.529 in the up conformation and complexed with ACE2 (PDB 7wpa chain A, at 2.77 Å [2], B.A.2 in the exclusively down conformation (PDB 7ub0, chain A, at 3.31 Å [3]), B.A.4/5 in the exclusively down conformation (PDB 7xnq, chain A, at 3.52 Å [4], and B.A.4/5 in the up conformation and complexed with ACE2 (PDB 7xwa, chain B, at 3.36 Å [5]. All structures were determined by cryoEM apart from PDB 7xwa, which was determined by X-ray crystallography.

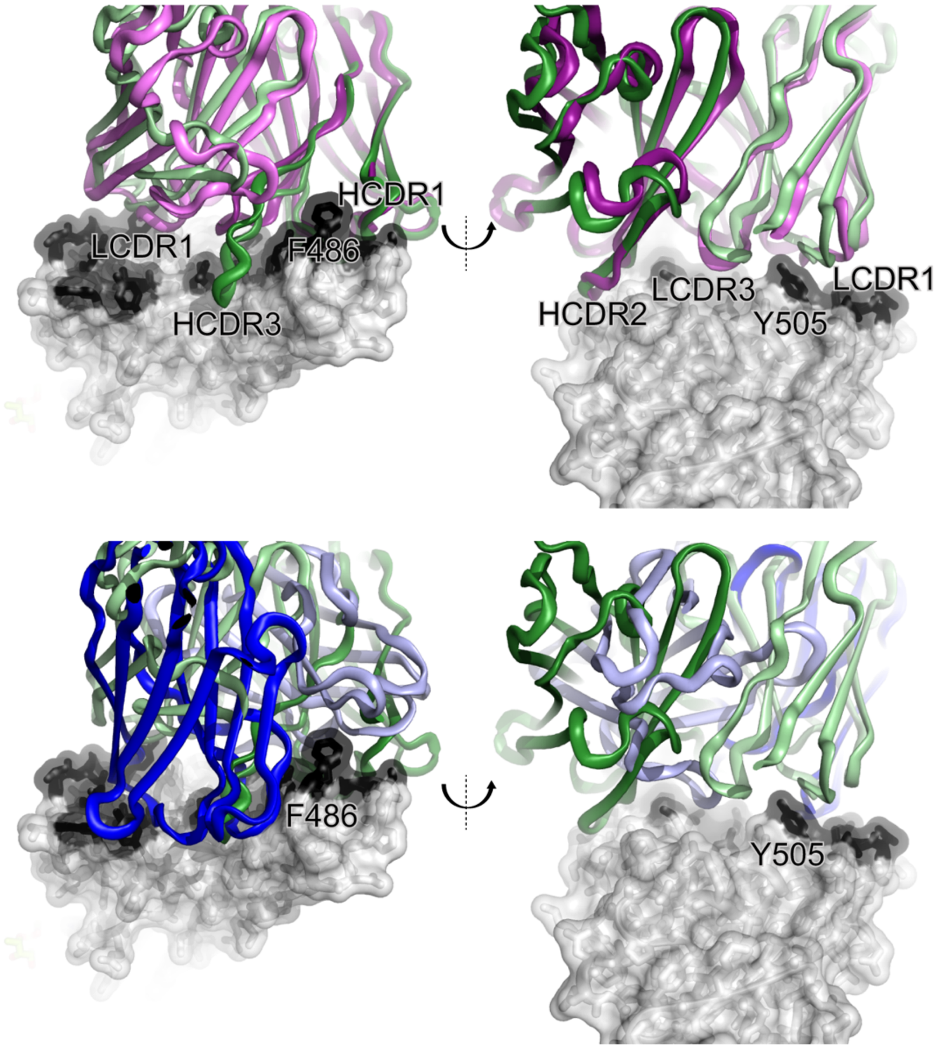


**Supplementary Fig. 8: Structural comparison of broadly neutralizing antibodies binding to the class 1/2 epitope (ACE-2 mimetics).** GAR05 (heavy and light chains as dark and light green cartoon) bound to the RBD (grey transparent surface with ACE2 interface coloured black) superposed with antibodies P2C-1F11 (top panels; dark and light purple for heavy and light chains) and S2K146 (bottom panels; dark and light blue for heavy and light chains). Prominent aromatic residues F486 and Y505 are indicated for reference.

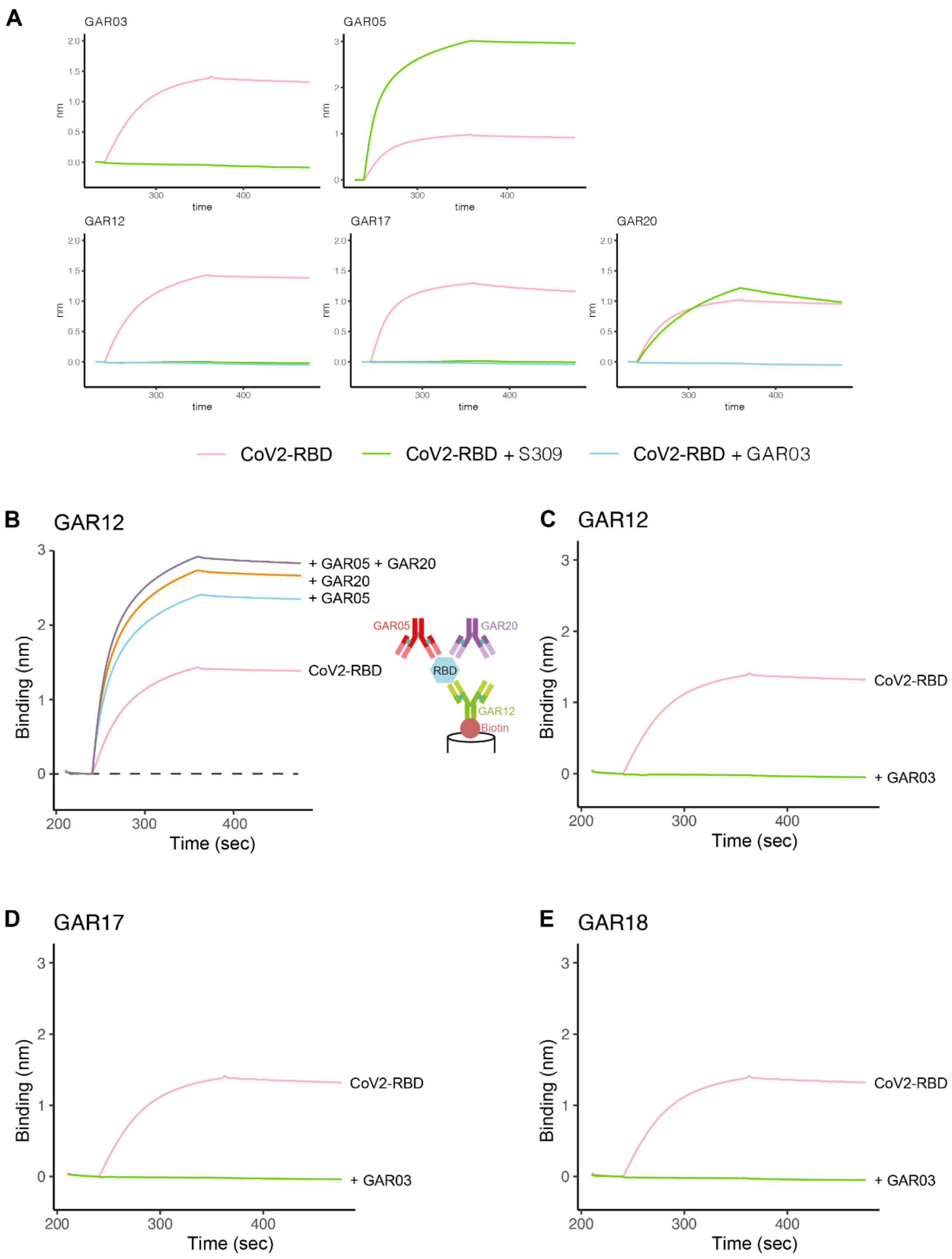


**Supplementary Fig. 9: S309 IgG, and GAR-antibody competition.** A: To assess antibody epitopes, biotinylated antibodies were captured onto streptavidin sensors and incubated with RBD (500 nM) pre-incubated with S309 IgG (1 µM) or GAR03 IgG (1 µM). B: BLI assay demonstrating the non-overlapping binding of GAR05, GAR12 and GAR20 on the surface of the RBD. Biotinylated GAR12 was captured onto streptavidin sensors and incubated with

RBD, RBD pre-incubated with GAR05, RBD pre-incubated with GAR20 or RBD pre-incubated with GAR05 and GAR20. Increased binding curves indicate that the antibodies do not compete.

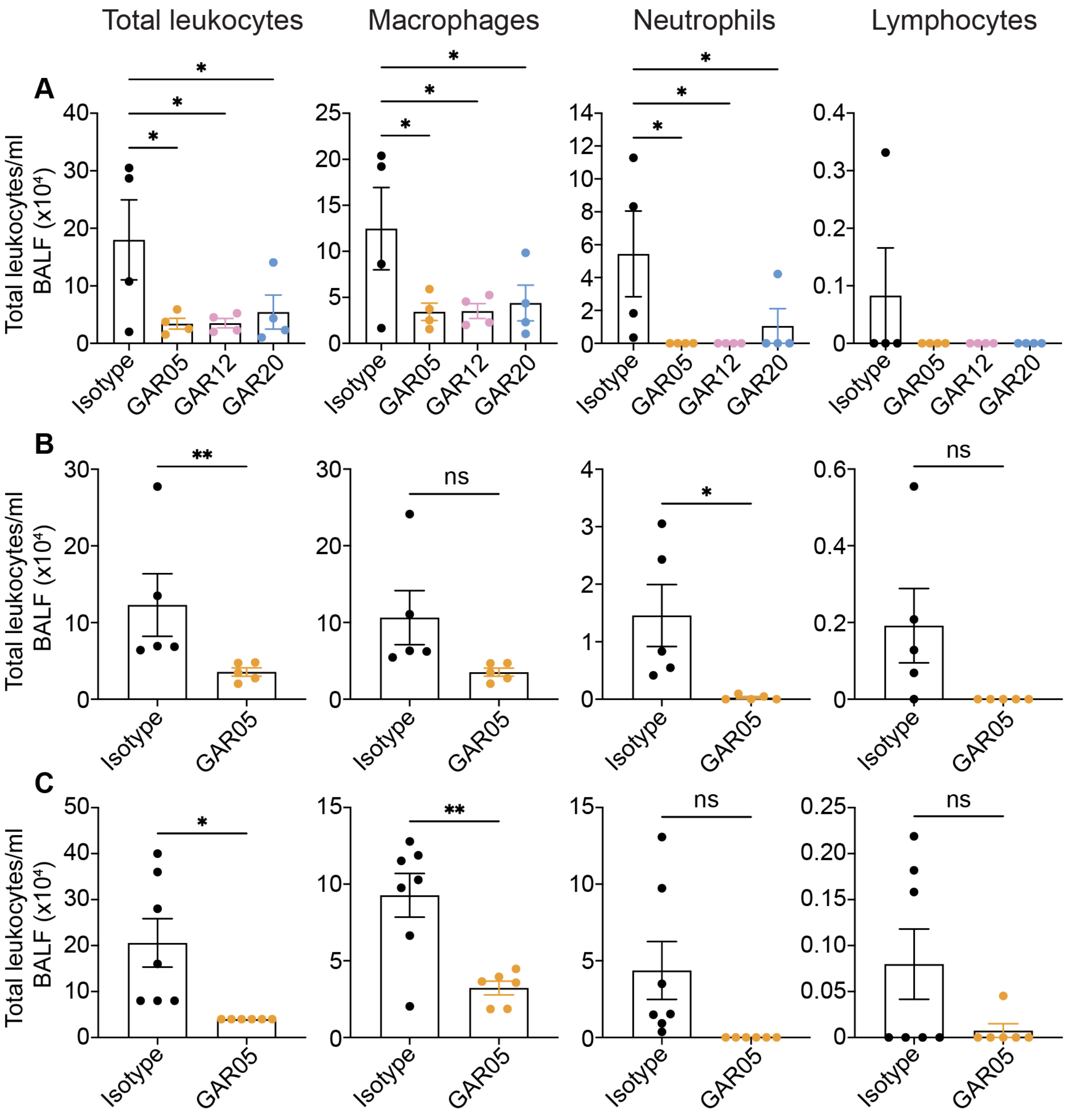


**Supplementary Fig. 10: BALF Differential cell counts from *in vivo* studies.** Differential leukocytes (macrophages, neutrophils and lymphocytes) cell counts isolated from bronchoalveolar lavage fluid (BALF) of Wuhan SARS-CoV-2. A) prophylaxis, and (B) therapeutic model, or (C) Delta SARS-CoV-2 prophylactic model.

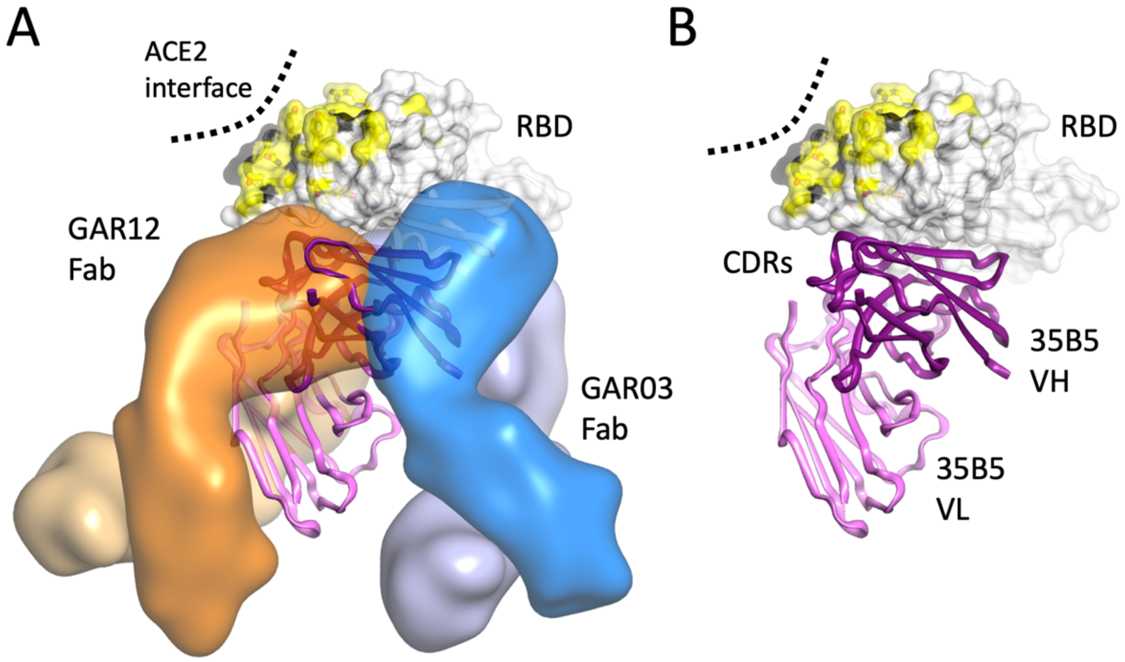


**Supplementary Fig. 11: Superposition of GAR03 and GAR12 Fabs with Fv from 35B5**. A: GAR12 and GAR03 Fabs (heavy and light chains as light/dark orange/blue transparent surfaces) positioned against the RBD (cream molecular surface and cartoon) whereas the ACE2 interface and overlayed omicron mutation positions are shaded black and yellow, respectively. The 35B5 Fv is shown as dark purple and violet cartoons (for heavy and light chains). B: The 35B5 antibody binds in an unusual manner with considerable contacts involving heavy-chain framework regions outside of the CDRs.

**Supplementary Table 1: Cryo-EM data collection and refinement.**

| | SARS CoV-2 VFLIP spike trimer + GAR03 (EMDB 25699) (PDB 7T5O) | SARS CoV-2 VLIP spike trimer + GAR05 (EMDB 25700) |
|---|---|---|
| **Data collection and processing** | | |
| Magnification | 130,000 | 130,000 |
| Voltage (kV) | 300 | 300 |
| Electron exposure ($e^{-}$/Å$^{2}$) | 50 | 50 |
| Defocus range (μm) | 0.8-1.8 | 0.8-1.8 |
| Pixel size (Å) | 1.08 | 1.08 |
| Symmetry imposed | C1 | C1 |
| Initial particle images (no.) | 475,165 | 542,046 |
| Final particle images (no.) | 130,129 | 341,602 |
| Map resolution (Å) 0.143 FSC threshold | 3.39 | 3.27 |
| **Refinement** | | |
| Map sharpening *B* factor (Å$^{2}$) | -78.3 | -88.5 |
| Initial models used (PDB) | 6xlu | |
| Model resolution (Å) 0.5 FSC threshold, Masked/unmasked | 7.6/7.8 | |
| *B* factor (Å$^{2}$) | | |
| Protein | 199.28 | |
| R.m.s deviations | | |
| Bond lengths (Å) | 0.007 | |
| Bond angles (°) | 1.384 | |
| Validation | | |
| MolProbity score | 1.5 | |
| Clashscore | 3.59 | |
| Poor rotamers (%) | 0.00 | |
| Ramachandran plot | | |
| Favored (%) | 94.95 | |
| Allowed (%) | 5.05 | |
| Disallowed (%) | 0.00 | |

**Supplementary Table 2: Diffraction data and model refinement statistics.** Values in parentheses represent values for the highest resolution shell.

| **X-Ray Diffraction Data Collection Statistics** | | | |
|---|---|---|---|
| Crystal | **GAR05 (CoV-2 RBD + GAR05 Fab)** | **GAR03 (CoV-2 RBD + GAR03 Fab + 10G4 Fab)** | **GAR12 (CoV-2RBD + GAR12 Fab)** |
| Wavelength | 0.9537 | 0.9537 | 0.9537 |
| Spacegroup | $P\ 3_2 2\ 2$ | $P\ 2_1 2_1 2_1$ | I 1 2 1 |
| Unit cell dimensions: a, b, c (Å); $\alpha, \beta, \gamma$, (°) | 111.29, 111.29, 112.86; *90.00, 90.00, 120.00* | 55.24, 102.08, 201.82; *90.00, 90.00, 120.00* | 93.61, 41.95, 183.03; *90.00, 97.08, 90.00* |
| Resolution range | 3.17-49.91 (3.18-3.40) | 2.73-49.48 | 2.25-48.08 (2.25-2.32) |
| Total reflections | 234263 (10010) | 372048 (39714) | 221099 (20451) |
| Unique reflections | 13965 (2418) | 31192 (3918) | 34125 (3150) |
| Completeness | 99.3 (96.5) | 99.4 (95.8) | 100.0 (100.0) |
| Multiplicity | 16.8 (16.6) | 11.9 (10.1) | 6.5 (6.5) |
| Average (I/σ(I)) | 21.4 (2.9) | 12.8 (1.7) | 17.0 (1.9) |
| Mean half set correlation, $CC_{1/2}$ | 0.998 (0.866) | 0.998 (0.672) | 0.999 (0.710) |
| Rmeas (all I+ and I-) | 0.079 (1.137) | 0.156 (1.384) | 0.060 (0.929) |
| Rpim (all I+ and I-) | 0.027 (0.392) | 0.047 (0.455) | 0.025 (0.394) |
| Wilson B (Å$^2$) | 126 | 54.5 | 48.7 |
| **Refinement and Model Statistics** | | | |
| $R_{work}/R_{free}$ | 0.237/0.308 | 0.223/0.269 | 0.211/0.254 |
| Complexes/asu | 1 | 1 | 1 |
| Additional Fab | none | Antibody 10G4* | none |
| Atoms protein | 4503 | 7883 | 4767 |
| B average protein (Å$^2$) | 134.4 | 64.13 | 72.06 |
| B average RBD(Å$^2$) | 134.6 | 53.51 | 54.4 |
| B average Heavy (Å$^2$) | 145.3 | GAR03: 68.32 10G4: 63.41 | 77.35 |
| B average Light (Å$^2$) | 123.8 | GAR03: 74.19 10G4: 61.14 | 84.29 |

| | | | |
|---|---|---|---|
| RMSD bond lengths (Å) | 0.003 | 0.003 | 0.009 |
| RMSD bond angles (°) | 0.809 | 0.706 | 0.951 |
| Ramachandran Outliers (%) | 0.49 | 0.86 | 0.33 |
| Ramachandran Favored (%) | 92.12 | 93.50 | 95.28 |
| **PDB entry** | **7t72** | **8dxu** | **8dxt** |

**Supplementary Table 3. GAR antibodies described in the study.**

| Antibody name | VH/VL germlines | VH CDR3 length (Kabat numbering) | Germline mutations |
|---|---|---|---|
| **GAR01** | IGHV3-30 & IGKV1-33 | 13 | 6 in VH, 1 in VL |
| **GAR03** | IGHV1-8 & IGLV3-21 | 12 | 4 in VH, 4 in VL |
| **GAR04** | IGHV1-2 & IGKV4-1 | 23 | 2 in VH, 1 in VL |
| **GAR05** | IGHV3-66 & IGKV1-33 | 14 | 8 in VH, 3 in VL |
| **GAR06** | IGHV2-70 & IGLV2-11 | 12 | 2 in VH, 4 in VL |
| **GAR07** | IGHV3-66 & IGKV1-33 | 9 | 4 in VH, 1 in VL |
| **GAR09** | IGHV3-66 & IGKV1-33 | 8 | 3 in VH, 2 in VL |
| **GAR11** | IGHV1-69 & IGLV2-14 | 17 | 10 in VH, 2 in VL |
| **GAR12** | IGHV3-23 & IGKV1-5 | 15 | 6 in VH, 1 in VL |
| **GAR13** | IGHV3-13 & IGKV1-39 | 15 | 4 in VH, 1 in VL |
| **GAR14** | IGHV3-13 & IGKV1-39 | 17 | 5 in VH, 4 in VL |
| **GAR15** | IGHV3-53 & IGKV1-9 | 10 | 3 in VH, 4 in VL |
| **GAR16** | IGHV3-13 & IGKV1-39 | 21 | 4 in VH, 4 in VL |
| **GAR17** | IGHV3-23 & IGLV1-47 | 17 | 6 in VH, 5 in VL |
| **GAR18** | IGHV4-59 & IGKV3-11 | 15 | 4 in VH, 2 in VL |
| **GAR20** | IGHV3-34 & IGLV3-21 | 12 | 6 in VH, 3 in VL |

1 Ye, G., Liu, B. & Li, F. Cryo-EM structure of a SARS-CoV-2 omicron spike protein ectodomain. *Nature Communications* **13**, 1214 (2022). https://doi.org:10.1038/s41467-022-28882-9
2 Yin, W. *et al.* Structures of the Omicron spike trimer with ACE2 and an anti-Omicron antibody. *Science* **375**, 1048-1053 (2022). https://doi.org:doi:10.1126/science.abn8863
3 Stalls, V. *et al.* Cryo-EM structures of SARS-CoV-2 Omicron BA.2 spike. *Cell Reports* **39**, 111009 (2022). https://doi.org:https://doi.org/10.1016/j.celrep.2022.111009
4 Cao, Y. *et al.* BA.2.12.1, BA.4 and BA.5 escape antibodies elicited by Omicron infection. *Nature* **608**, 593-602 (2022). https://doi.org:10.1038/s41586-022-04980-y
5 Kimura, I. *et al.* Virological characteristics of the SARS-CoV-2 Omicron BA.2 subvariants, including BA.4 and BA.5. *Cell* **185**, 3992-4007.e3916 (2022). https://doi.org:https://doi.org/10.1016/j.cell.2022.09.018
